# Two species of the *Entoloma quadratum*–*murrayi* complex from Japan, *E. kermesinum* sp. nov. and *E. flavescens* sp. nov.

**DOI:** 10.1101/2024.04.11.589125

**Authors:** Hirotoshi Sato, Otomi Satomi, Yoriko Sugiyama

## Abstract

We describe two new species, *Entoloma. kermesinum* sp. nov. and *E. flavescens* sp. nov., which are confused with *E. quadratum* and *E. murrayi*, respectively. We sequenced the large subunit of mitochondrial ribosomal RNA, the nuclear ribosomal internal transcribed spacer region and 22 single-copy genes for 51 specimens of *E. kermesinum*, *E. flavescens*, *E. album*, and related species. Species boundaries were assessed using the molecular phylogenetics and population genetics approaches. Specimens of *E. kermesinum*, *E. flavescens*, and *E. album* formed independent clades, which were phylogenetically distinct from the specimens of *E. quadratum* and *E. murrayi* collected around the type locality (i.e., New England). Although the phylogenetic distance between *E. flavescens* and *E. album* was small, gene flow between them was restricted in areas where they coexisted, suggesting reproductive isolation. Therefore, these five species can be treated as independent species. We found characteristics useful for identifying *E. kermesinum* and *E. flavescens*. In particular, *E. kermesinum* is characterized by a crimson to brown-red and fibrillose pileus, finely covered by whitish fibrous scales; *E. flavescens* is characterized by a lemon-yellow to tan and shiny-to-silky pileus. In addition, relatively large basidiospores and clamp connections are diagnostic features of these two species.

## Introduction

Species delimitation is one of the most important challenges in fungal classification. Because fungi have simple and plastic morphological characteristics, cryptic species are often confused even in macrofungi that form conspicuous fruitbodies [1, 2]. By contrast, known species may have many synonyms in fungal taxa. Indeed, a recent study estimated the total number of operational taxonomic units (OTUs) for all macrofungal taxa by examining the increase in richness of operational taxonomic units (OTUs) with increasing the number of accessions in the nucleotide sequence database, suggesting that the number of species in some fungal taxa, such as *Agaricus* and *Marasmius*, may have been overestimated due to occurrence of synonymous species [3]. Consequently, taxonomic studies that use DNA-based approaches, such as sequencing of fungal barcodes [4, 5], comparative gene genealogies [6–8], and population genetics [2, 9, 10], are needed for species delimitation of many fungal taxa.

*Entoloma* (Entolomataceae, Agaricales) is the type-genus of the family Entolomataceae and one of the most abundant and ubiquitous macrofungal taxa, almost worldwide [11–19]. *Entoloma* is characterized by a pink to salmon-pink spore print and angular basidiospores [20, 21]. Molecular phylogenetic studies have shown that *Entoloma* is a monophyletic group, sister to genera *Clitopilus* and *Clitocella* [3, 22, 23]. Most species of this genus are soil and/or litter saprotrophs [24, 25]. However, some species, such as *E. sarcopum* Nagas. & Hongo, *E. sinuatum* (Bull.) P. Kumm. and *E. rhodopolium* (Fr.) P. Kumm., are ectomycorrhizal, forming a mutualistic relationship with host plants [26], while *E. clypeatum* forms ectomycorrhiza-like roots with Rosaceae and Ulmaceae [27]. Notably, many new *Entoloma* species have been discovered in recent years (Index Fungorum), and thus undescribed species are undoubtedly present in this genus. Nevertheless, the number of described *Entoloma* species (i.e., 2,603; the Index Fungorum) is much larger than their estimated global OTU richness (i.e., 698 [3]), suggesting that not only many undescribed species, but also many synonymous species are present in *Entoloma*. Overall, the taxonomy of this genus must be resolved.

In Japan, more than 50 *Entoloma* species are known (Imazeki & Hongo 1987), but the taxonomy of this genus remains to be resolved. For instance, *E. quadratum* (Berk. & M.A. Curtis) E. Horak, *E. murrayi* (Berk. & M.A. Curtis) Sacc. & P. Syd., and *E. album* Hiroë are morphologically similar, except that *E. quadratum*, *E. murrayi* and *E. album* are characterized by reddish, yellowish and whitish basidiocarps, respectively [28]. However, basidiocarps with intermediate colors (e.g., tan and orange-tan) are often found; therefore, it remains unclear whether these three species should be treated as distinct species. Among these three species, the type localities of *E. quadratum* and *murrayi* are the Eastern United Sates (i.e., New England), whereas that of *E. album* is Japan. In the type locality of *E. quadratum* and *E. murrayi*, the former species is distinguished from the latter species in having clamp connections [11], but this feature has not been confirmed for Japanese specimens. Moreover, no DNA-based study has confirmed whether the Japanese specimens are conspecific to their North American counterparts. Therefore, besides detailed morphological examination, DNA-based approaches are needed to clarify the taxonomy of the *E. quadratum*– *murrayi* complex observed in Japan.

This study aimed to taxonomically reconsider fungal materials known as *E. quadratum* (Japanese name: Aka-ibokasatake), *E. murrayi* (Japanese name: Ki-ibokasatake) and *E. album* (Japanese name: Shiro-ibokasatake) in Japan. Molecular phylogeny and population genetics approaches showed that *E. quadratum*, *E. murrayi*, and *E. album* are distinct species. Furthermore, our results indicated that fungal materials known as *E. quadratum* and *E. murrayi* in Japan should be treated as new species (hereafter, *E. kermesinum* and *E. flavescens*, respectively). Here, we discuss the taxonomy of *E. kermesinum*, *E. flavescens* and *E. album*.

## Materials & Methods

### Field survey

From Sep 2018 to Jul 2021, 51 specimens of *E. kermesinum*, *E. flavescens*, *E. album*, and related species were collected from eight forest sites in Japan (S1 Table). All necessary permits were obtained for the described field study, which complied with all relevant regulations. Small sections of fruit bodies were removed and stored in 99.5% ethanol for molecular analysis, and remaining sections were dried and preserved as voucher specimens. Dried specimens were deposited in the National Museum of Nature and Science, Tokyo (TNS) and the herbarium of Kyoto University (KYO). Six samples used in other study (i.e., samples collected in Yakushima, Kagoshima Pref., Japan) [29] were included in the analysis.

### DNA extraction, PCR amplification, and sequencing

Total DNA was extracted from tissues of voucher specimens using the cetyltrimethylammonium bromide (CTAB) method, as described previously [10]. Two-step PCR was performed to amplify the following target regions: the large subunit of mitochondrial ribosomal RNA (mtLSU), the nuclear ribosomal internal transcribed spacer (ITS) region, and 40 single-copy genes used in other studies [30, 2020]. In the first step, fragments of the target regions were amplified using predesigned primers [30, 2020, 31] listed in S2 Table. The primers contain Illumina sequencing primer regions and 6-mer Ns (forward: 5’-TCGTCGGCAGCGTCAGATGTGTATAAGAGACAGNNNNNN [specific primer]-3’ and reverse: 5’-GTCTCGTGGGCTCGGAGATGTGTATAAGAGACAGNNNNNN [specific primer]-3’). Illumina sequencing adaptors plus 8-bp identifier indices [32] were added in the subsequent PCR using a forward and reverse fusion primer (forward: 5’-AATGATACGGCGACCACCGAGATCTACAC-index-TCGTCGGCAGCGTC-3’ and reverse: 5’-CAAGCAGAAGACGGCATACGAGAT-index-GTCTCGTGGGCTCGG-3’). The PCR conditions of first and second steps were as described previously [30]. An equal volume of the respective PCR products was pooled, and 450–600 bp amplicons were excised and extracted with the E-Gel SizeSelect 2% agarose gel system (Thermo Fisher Scientific). Amplicon libraries were sequenced by 2 × 250 bp paired-end sequencing on a MiSeq platform (Illumina, San Diego, CA, USA) using MiSeq v2 Reagent NANO Kit, according to the manufacturer’s instructions.

### Bioinformatics analysis

BCL2FASTQ version 1.8.4 (Illumina) was used to convert the base calls into forward, index1, index2, and reverse FASTQ files. The FASTQ files were demultiplexed using “clsplitseq” command in Claident ver. 0.2.2019.04.27 [33]. In total, 4,528,951 demultiplexed reads were deposited in the DDBJ Sequence Read Archive (DRA accession: DRA017274, DRA017275, and DRA01276). Index reads and forward and reverse primer positions were trimmed during this step. VSEARCH software [34] was used to merge forward and reverse reads of sequences in each sample. Merged sequence reads with low quality scores (<Q30) were removed with “clfilterseq” command in Claident. Reads likely to include a high proportion of sequencing errors (noisy reads) were eliminated using “clcleanseq” command of Claident with default setting. Chimeric sequences were eliminated with a minimum score of 0.1 to report chimeras using UCHIME [35]. The processed reads were assembled into contigs (unique sequences) in Claident using a similarity cut-off of 100%. The final Claident output files were processed using R version 4.3.0 [36], as described previously [2]. Unique sequences of each gene were separately aligned with the nucleotide sequences of the same gene in *P. chrysosporium* using MAFFT v7.24559 [37]. Gene sequences that were not detected in >70% of the total samples were removed. Alignment data of the remaining genes (FG1024, FG459, FG465, FG507, FG525, FG533, FG534, FG576, FG579, FG588, FG595, FG756, FG761, FG771, FG927, FG975, MS320, MS353, MS358, MS361, MS378, MS453, ITS1, ITS2, and mtLSU) were analyzed based on population genetics and molecular phylogenetic inference.

### Molecular Phylogenetic inference

Sequences obtained from the same sample and locus were incorporated into the consensus sequence (IUPAC standard) using the “consensus” function in “seqinr ver. 3.4-5” in R package [38]. The consensus sequences were used for the subsequent molecular phylogenetic inference. GenBank accession numbers are provided for the consensus sequences of ITS1 and ITS2 regions (S1 Table). Molecular phylogenetic inference was performed using the mtLSU, ITS, and the nuc_concat datasets (the concatenated sequences of 22 single-copy genes) separately. Because one locus (MS444) was incongruent with several other loci of the nuc_concat dataset based on the congruence among distance matrices (CADM) test, the remaining 21 loci were subsequently analyzed.

Phylogenetic inference based on the maximum likelihood (ML) method was performed using RAxML ver. 8.1.5 [39], in which tree searches were repeated 15 times using random sequence addition for generating starting trees. Bootstrap support values were calculated from 1,000 standard bootstrap replications in RAxML. Parameters of the GTR Gamma model were estimated separately for each partition according to model selection, based on the Akaike information criterion using Kakusan 4 [40]. The input and output data are deposited in Dryad (doi: 10.5061/dryad.pg4f4qrws).

### Population genetics analysis

Using “as.matrix.alignment” function in R package “seqinr”, the alignment of each nuclear locus was converted into a matrix of genotypes, where rows and columns represented samples and nucleotide positions, respectively. To reduce biases, samples and loci with many missing datapoints were removed (i.e., 25% cut-off levels). The data frame was converted into a genind object, and AMOVA was performed to determine the proportion of nuclear genetic variation that could be attributed to differences between OTUs, between geographical regions (sampling localities), and between or within samples using “poppr.amova” function in R package “poppr ver. 2.8.3” [41]. This test also calculated *Φ* statistics, analogous to Wright’s *F*-statistics. The statistical significance of variance components was computed using a permutation test (9,999 iterations) implemented in R package “pegas ver. 1.2” [42].

### Observation of morphological characteristics

The macro- and micromorphological characteristics of basidiomes were described from fresh and dried specimens, respectively. Microscopic observations were performed under a CX40 phase contrast microscope (OLYMPUS, Tokyo) with sections of basidiome tissue mounted in 3% potassium hydroxide. Basidiospore measurements were taken at 1,000× magnification under an optical microscope. Each color was described using CMYK (cyan, magenta, yellow, and key plate) color chart. The lengths and widths of 10 basidiospores, 5 cheilocystidia and 3–5 basidia were measured for each collection. Between-group differences in lengths and widths were analyzed with the Steel-Dwass test using R.

### Nomenclature

The electronic version of this article in Portable Document Format (PDF) in a work with an ISSN or ISBN will represent a published work according to the International Code of Nomenclature for algae, fungi, and plants, and hence the new names contained in the electronic publication of a PLOS article are effectively published under that Code from the electronic edition alone, so there is no longer any need to provide printed copies.

The new names contained in this work have been submitted to MycoBank, where they will be made available to the Global Names Index. The unique MycoBank number can be resolved, and the associated information can be viewed through any standard web browser by appending the MycoBank number contained in this publication to the prefix http://www.mycobank.org/MB/. The online version of this work was archived and made available in the following digital repositories: PubMed Central and LOCKSS.

## Results

### Molecular phylogenetic inference

The mtLSU dataset consisted of 283 nucleotide sites, of which 66 were variable. For the mtLSU dataset, both *E. kermesinum* and *E. album* had single haplotypes, whereas *E. flavescens* had three haplotypes. The ML tree of mtLSU dataset showed that *E. kermesinum* was separated from *E. album* and *E. flavescens*, but *E. album* and *E. flavescens* were not clearly separated from each other (Fig. 1A).

**Fig. 1.**
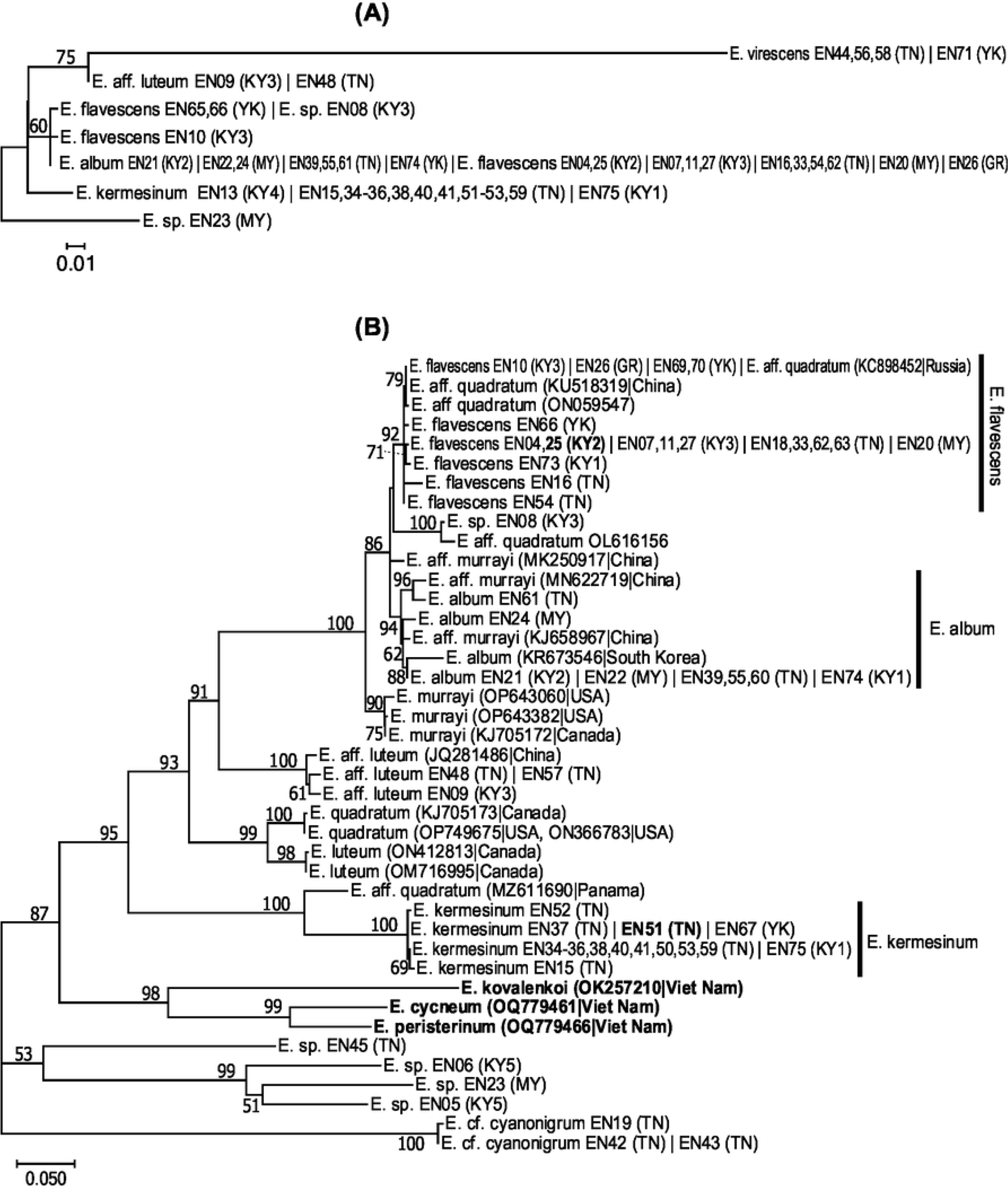
Maximum-likelihood (ML) trees of the *Entoloma kermesinum*, *E. flavescens*, *E. album* and the related species, inferred from the nucleotide sequences of (A) the large subunit of mitochondrial ribosomal RNA (mtLSU) and (B) nuclear ribosomal internal transcribed spacer (ITS) region. Numbers near the branches are bootstrap values (>50%). Samples with identical sequences are premerged as unique sequences. Each taxon name is followed by strain name and locality abbreviation—Strain name (Locality abbreviation). Bold indicates a holotype specimen. Locality abbreviations are as follows: Kutashimonocho, Kyoto, Japan (KY1), Yoshidayama, Kyoto, Japan (KY2), Mt. Daimonji, Kyoto, Japan (KY3), Sasari-Toge Pass, Kyoto, Japan (KY4), Fushimi-inari, Kyoto, Japan (KY5), Tanakami, Shiga, Japan (TN), Mt. Maya, Hyogo, Japan (MY), Gero, Gifu, Japan (GR), and Yakushima, Kagoshima, Japan (YK).

The nuclear ITS dataset consisted of 728 nucleotide sites (ITS1: 342 bp, ITS2: 386 bp), of which 399 were variable. The ML tree of the ITS dataset showed that specimens of *E. kermesinum* formed a monophyletic clade with high bootstrap support (BS = 100%) (Fig. 1B). Maximum likelihood distances within *E. kermesinum* were substantially small (0–0.0038). The *E. kermesinum* clade was phylogenetically distinct from the clades of other species. Notably, the *E. kermesinum* clade was paraphyletic to the *E. quadratum* clade, which comprised specimens collected near the type locality of *E. quadratum* (i.e., New England). The *E. kermesinum* clade was sister to the specimens of *E.* aff. *quadratum* collected in Panama (MZ611690) with 100% BS, but ML distances were high (0.106– 0.115). The *E. quadratum* clade was sister to that of *E. luteum*, collected in Canada. Specimens of *E. album* collected in Japan (this study) and South Korea (KR673546) and specimens of *E.* aff. *murrayi* collected in China (KJ658967 and MN622719) formed a monophyletic clade (BS = 94%), although the South Korean specimen had a somewhat distinct sequence (the ML distances from other specimens ranged from 0.041 to 0.055). The ML distances within the *E. album* clade, excluding the South Korean specimen, ranged from 0 to 0.030. Specimens of *E. flavescens* and several specimens from other countries, including *E.* aff. *quadratum* collected from China (KU518319) and Russia (KC898452), formed a monophyletic clade (BS = 92%). The ML distances within the *E. flavescens* clade ranged from 0 to 0.019. The *E. flavescens* clade was closely related to the *E. album* clade, rather than the *E. murrayi* clade (BS = 86%). The ML distances between *E. flavescens* and *E. album* ranged from 0.011 to 0.051, excluding the South Korean *E. album* specimen, whereas those between *E. flavescens* and *E. murrayi* ranged from 0.030 to 0.056. One specimen of *E.* sp. (EN08) was somewhat closely related to *E.* aff. *quadratum* (OL616156) (ML distance = 0.012), and both formed a monophyletic clade (BS = 100%). This clade was somewhat distinct from both of the *E. album* clade (ML distance = 0.054–0.081, excluding the South Korean specimen) and the *E. flavescens* clade (ML distance = 0.039–0.075). The clade comprising *E. album*, *E. flavescens*, *Entoloma* sp. (EN08), and *E. murrayi* was closely related to that of the Japanese specimens of *E.* aff. *luteum* and the North American specimens of *E. quadratum* and *E. luteum* (BS = 93%), rather than *E. kermesinum*. Moreover, three species that were recently reported from Viet Nam (i.e., *E. kovalenkoi*, *E. cycneum*, and *E. peristerinum*) were phylogenetically distinct from other species.

The nuc_concat dataset consisted of 7,432 nucleotide sites, of which 2,820 were variable. Specimens of *E. kermesinum* formed a monophyletic clade (BS = 100%) that was phylogenetically distinct from other species (Fig. 2). Maximum likelihood distances within the *E. kermesinum* clade were small (0–0.006). Specimens of *E. album* clustered into a monophyletic clade (BS = 100%), and the ML distance within the clade ranged from 0.000 to 0.008. The monophyly of the *E. flavescens* specimens was strongly supported (BS = 100%). Maximum likelihood distances within the *E. flavescens* clade were somewhat larger than those in other clades, especially between specimens collected in Yakushima and other areas (0–0.013). Notably, *E. album*, *E. flavescens* and *E*. sp. (EN08) were closely related. The *E. album* clade was sister to the *E. flavescens* clade (BS = 100%) (ML distances = 0.017–0.029). The clade comprising *E. album* and *E. flavescens* was sister to E. sp. (EN08), and the ML distances ranged from 0.042 to 0.056. The clade of *E. album*, *E. flavescens* and *E*. sp. (EN08) was sister to the clade of *E.* aff. *luteum*, but ML distances were large (0.138–0.201).

**Fig. 2.**
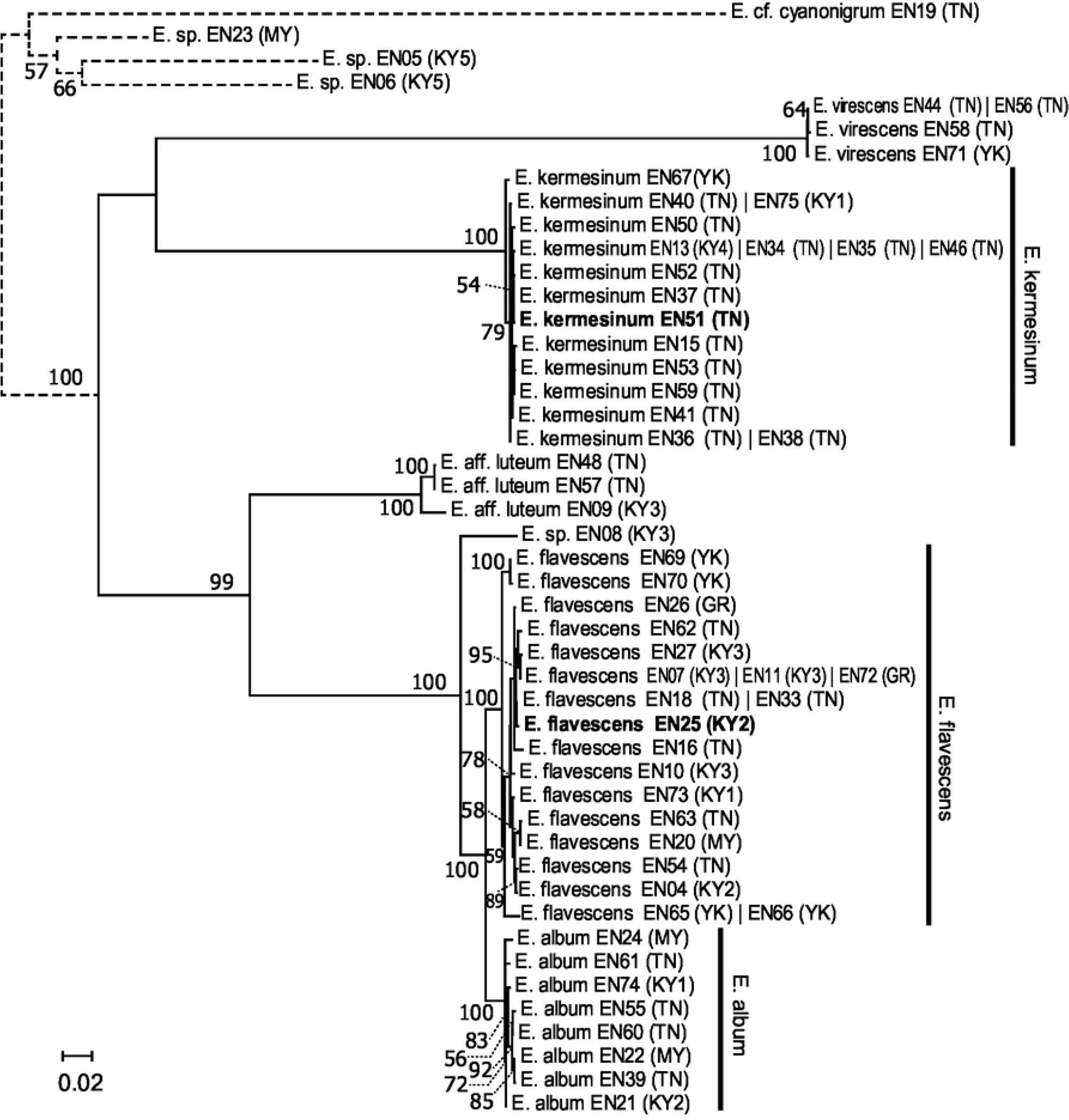
Maximum-likelihood (ML) trees of the *Entoloma kermesinum*, *E. flavescens*, *E. album* and the related species, inferred from the nucleotide sequences of concatenated 21 nuclear single copy genes. Numbers near the branches are bootstrap values (>50%). Samples with identical sequences are premerged as unique sequences. Each taxon name is followed by strain name and locality abbreviation—Strain name (Locality abbreviation). Bold indicates a holotype specimen. Locality abbreviations are as follows: Kutashimonocho, Kyoto, Japan (KY1), Yoshidayama, Kyoto, Japan (KY2), Mt. Daimonji, Kyoto, Japan (KY3), Sasari-Toge Pass, Kyoto, Japan (KY4), Fushimi-inari, Kyoto, Japan (KY5), Tanakami, Shiga, Japan (TN), Mt. Maya, Hyogo, Japan (MY), Gero, Gifu, Japan (GR), and Yakushima, Kagoshima, Japan (YK).

### Population genetics analysis

Results of AMOVA indicated significant and large genetic differentiation in the nuc_concat dataset between *E. album* and *E. flavescens*. The genetic differentiation between *E. album* and *E. flavescens* explained 71.1% of total variance, and *Φ* value was significantly higher than 0 (*Φ* = 0.711, P = 0.0029; Table 1), indicating highly restricted gene flow between these two species. In addition, variation was explained by differentiation among geographical regions within each species (12.9%, *Φ* = 0.445, P < 0.001) (Table 1), suggesting restricted gene flow among different geographical regions. The remaining variation was explained by differentiation between and within samples (16.1%). The results were almost identical when the cut-off levels of samples and loci were changed.

**Table 1.**
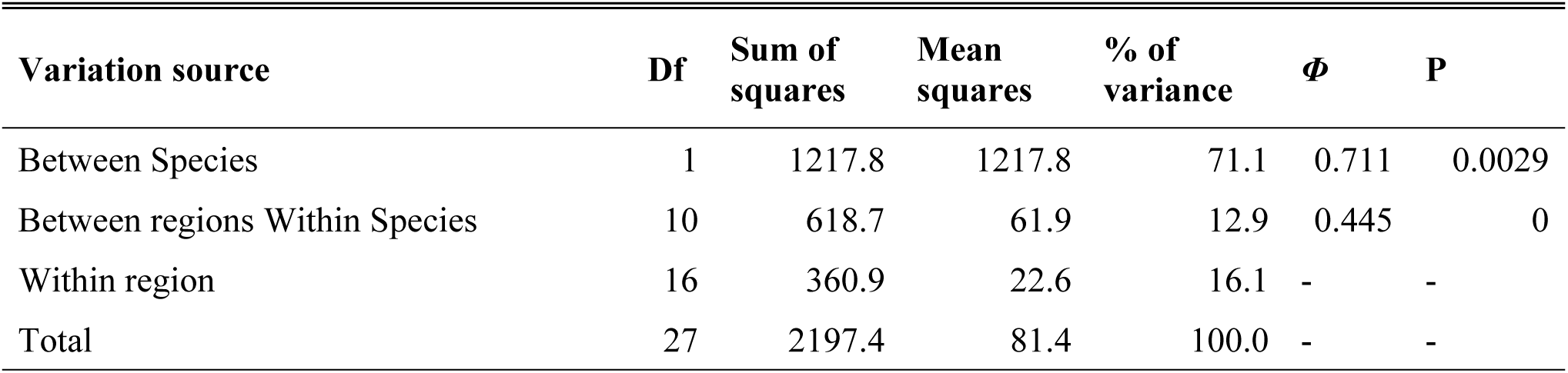
Results for analysis of molecular variance (AMOVA) based on single nucleotide polymorphism of nuclear loci, considering species and regions (locality). AMOVA is a hierarchical analysis of molecular variance, estimating the % of molecular variance accounted for by each level of the nested sampling hierarchy as well as *Φ* (≒*F*_st_).

| Variation source | Df | Sum of squares | Mean squares | % of variance | $\Phi$ | P |
| --- | --- | --- | --- | --- | --- | --- |
| Between Species | 1 | 1217.8 | 1217.8 | 71.1 | 0.711 | 0.0029 |
| Between regions Within Species | 10 | 618.7 | 61.9 | 12.9 | 0.445 | 0 |
| Within region | 16 | 360.9 | 22.6 | 16.1 | - | - |
| Total | 27 | 2197.4 | 81.4 | 100.0 | - | - |

### Observation of morphological characteristics

Differences in macroscopic (Fig. 3) and microscopic (Figs. 4–6) morphologies were found among the three species. Basidiocarps of *E. kermesinum* and *E. album* were characteristically crimson (Fig. 3A–C) and whitish colors (Fig. 3G–I), respectively. Most *E. flavescens* specimens had lemon-yellow to yellow basidiocarps, but some specimens (EN18, EN20 and EN70) were characterized by a tan-orange basidiocarp (Fig. 3D–F). Specimens of *E. kermesinum* had a fibrillose pileus covered by whitish fibrous scales, whereas *E. album* and *E. flavescens* had a shiny- to-silky pileus (Fig. 3A-I). Basidiospore length and width of *E. kermesinum* were significantly larger than those of *E. album* and *E. flavescens*, but the difference was small (Figs. 4–6). Moreover, cheilocystidia widths in *E. album* were significantly smaller than those in *E. flavescens* and *E. kermesinum* (Figs. 4–6).

**Fig. 3.**
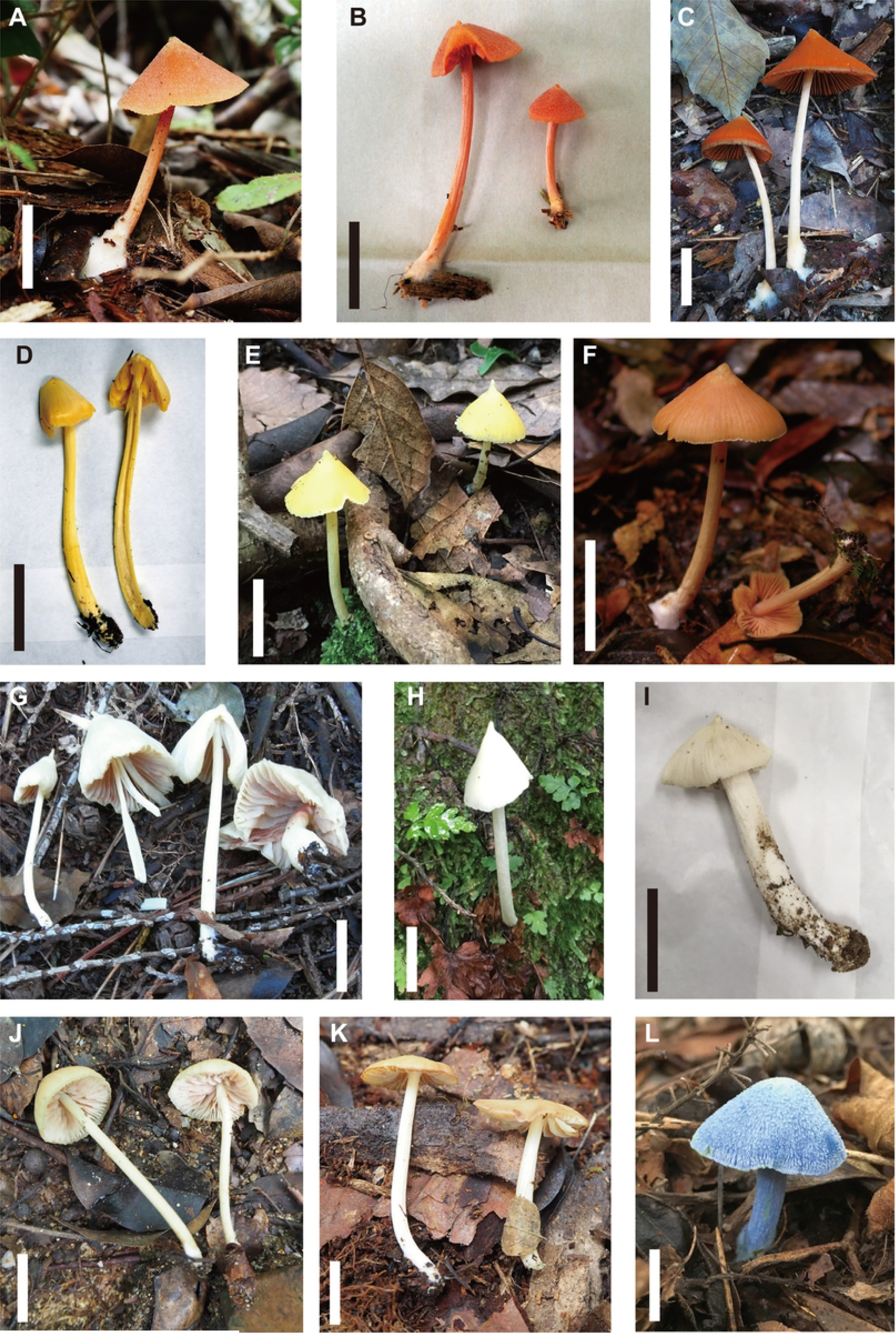
Photographs of the basidiomes of *Entoloma kermesinum*, *E. flavescens*, *E. album* and related species. (A) *E. kermesinum* (TNS-F-83013), (B) *E. kermesinum* (TNS-F-83014; holotype), (C) *E. kermesinum* (TNS-F-83002), (D) *E. flavescens* (TNS-F-82996), (E) *E. flavescens* (TNS-F-82995; holotype), (F) *E. flavescens* (KYO-YK489), (G) *E. album* (TNS-F-83023), (H) *E. album* (TNS-F-82291), (I) *E. album* (TNS-F-82292), (J) *E.* sp. (TNS-F-82981), (K) *E.* aff. *luteum* (TNS-F-82982), and (L) *E. virescens* (TNS-F-83019). *Bars*: 2 cm.

**Fig. 4.**
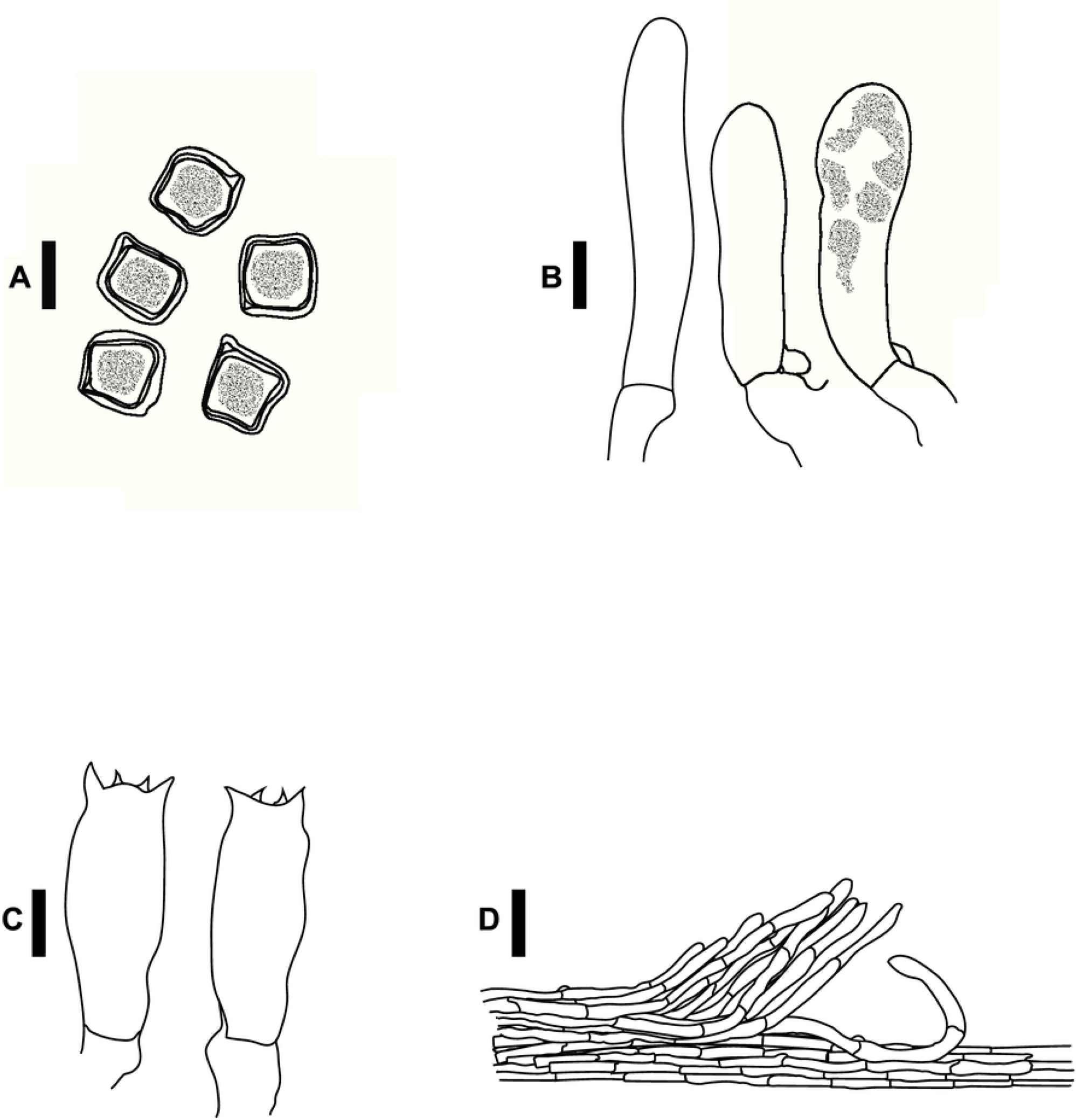
Basidiospores (A), cheilocystidia (B), basidia (C), and pileipellis hyphae (D) of *Entoloma kermesinum*. *Bars*: A–C 10 µm; D 50 µm.

**Fig. 5.**
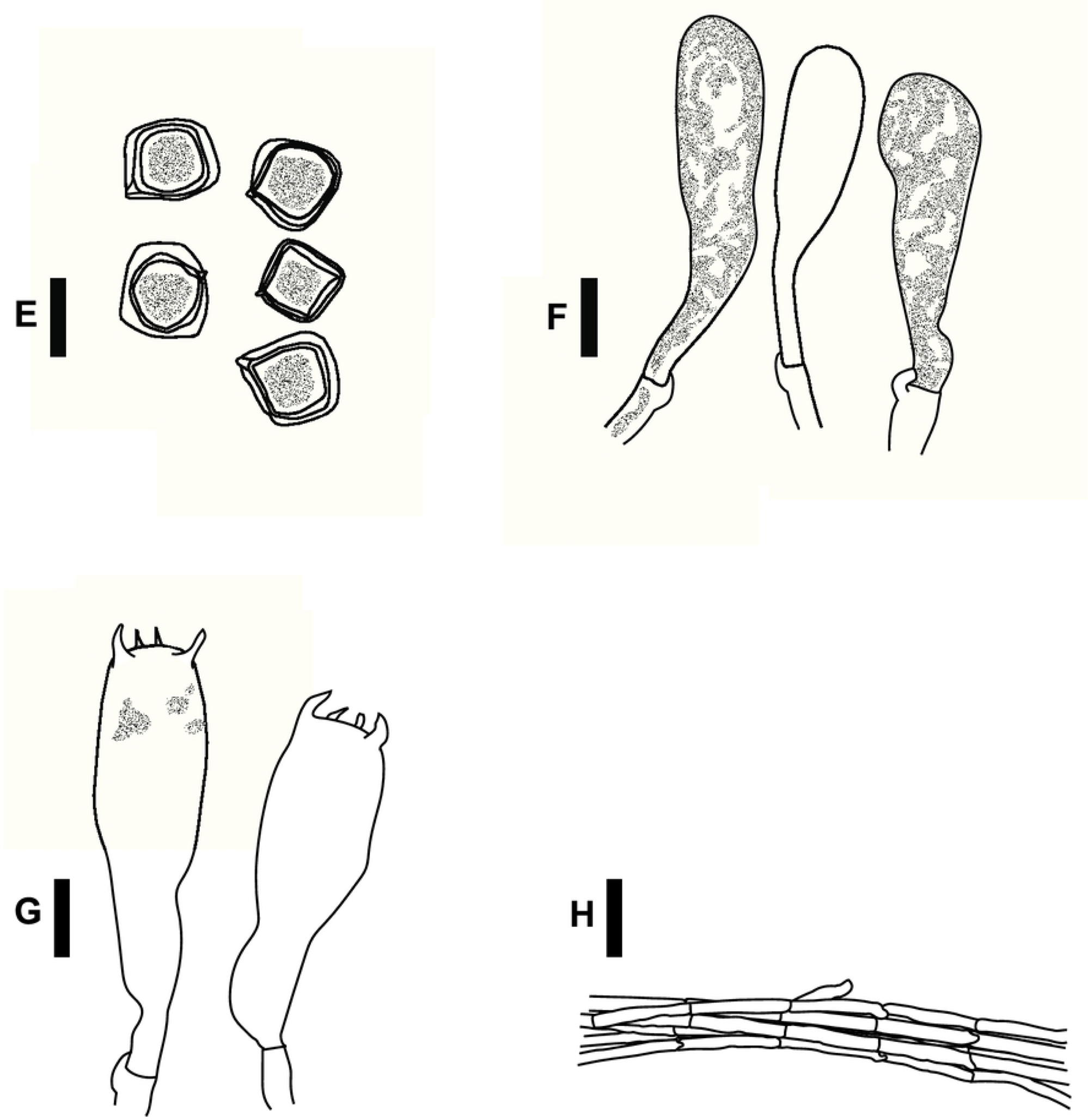
Basidiospores (A), cheilocystidia (B), basidia (C), and pileipellis hyphae (D) of *Entoloma flavescens*. *Bars*: A–C 10 µm; D 50 µm.

**Fig. 6.**
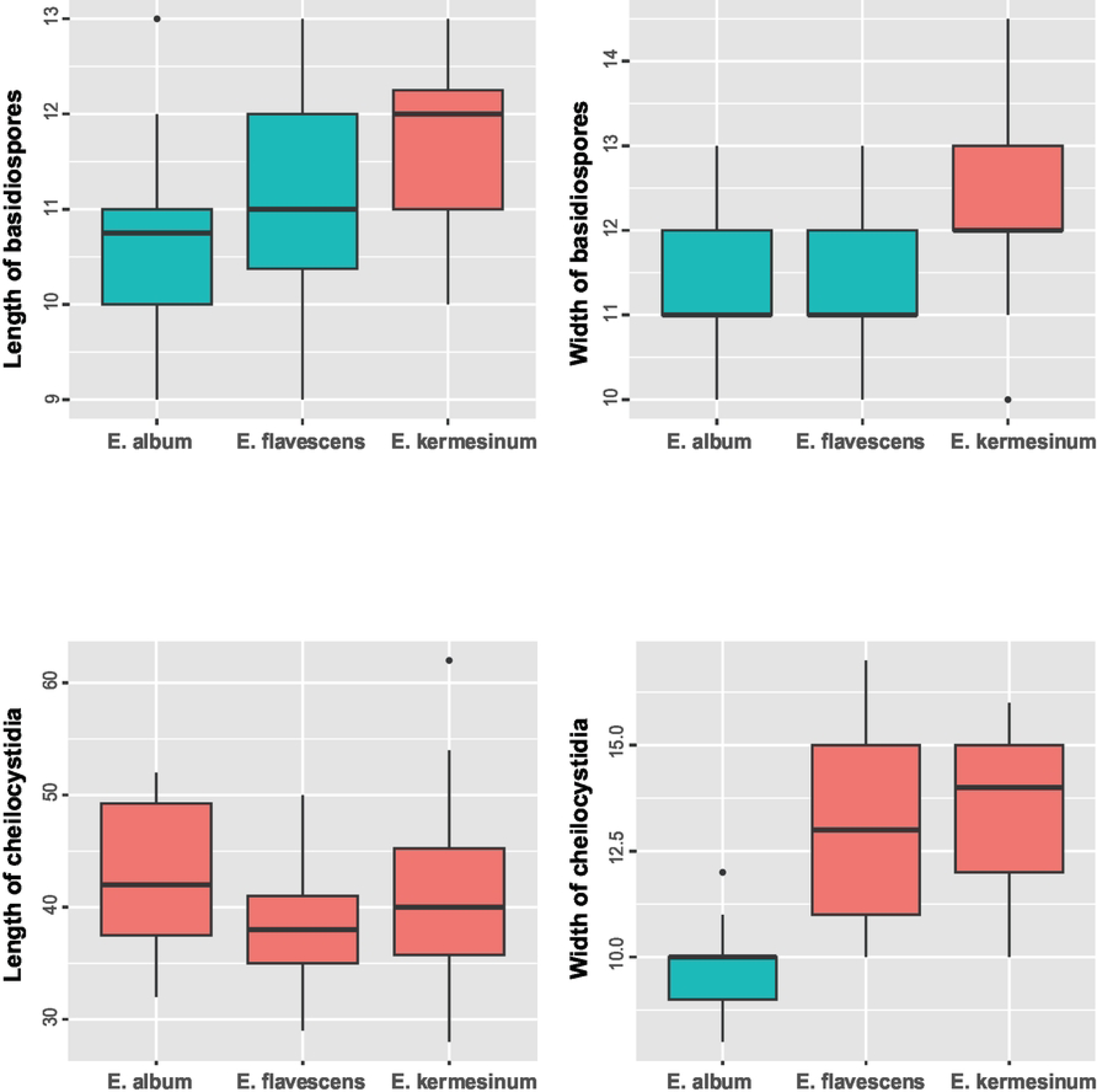
Differences in the width and length in basidiospores and cheilocystidia between *Entoloma kermesinum*, *E. flavescens* and *E. album*. Box plot displaying the median (bold horizontal line), first and third quartiles (“rectangle”), and 95% confidence intervals (“solid vertical line”). Different colors in box plots indicate significant differences in the Steel–Dwass test (α= 0.05).

### Taxonomy

#### *Entoloma kermesinum* Hirot. Sato, sp. nov. (Figs. 3A–C, 4)

##### MycoBank no

MB 10015983.

##### Holotype

Tanakami, Otsu, Shiga Pref., Japan, on soil in a mixed forest of *Quercus serrata* and *Pinus densiflora*, Sep. 30, 2019, Voucher (**TNS-F-83014**), GenBank accession numbers: ITS1 (LC786677), ITS2 (LC786729)

##### Etymology

*kermesinum*, Latin, crimson color

##### Diagnosis

*Entoloma kermesinum* sp. nov. is characterized by the following characteristics, a crimson to brown-red pileus, finely covered by whitish fibrous scales, especially in an acute cusp at the apex, basidiospores 10–13× 10–14.5 µm, pileipellis a cutis composed of cylindrical to subcylindrical hyphae with scattered bundles of ascending hyphae, and clamp connections in all hyphal tissues.

##### Description

Pileus 1–6 cm diam, thin, conical to campanulate, with an acute cusp at the apex; surface crimson (C 14, M92, Y100, K 0) to brown red (C 26, M 76, Y 96, K 0), deep red (C 33, M 93, Y 100, K 0), dry, densely fibrillose with whitish fibrous scales specifically near the apex, longitudinally striate except near the apex when wet, margin slightly lobed. Lamellae adnexed to almost free, subdistant, subventricose, subdenticulate on the edges, concolorous with pileus or slightly darker. Flesh thin, fragile, concolorous with pileus. Stipe 4–10 cm long, 2–4 mm thick, slender, equal or subbulbous, striate fibrillose, often twisted, hollow, concolorous with pileus or paler, sometimes whitish, base white-mycelioid. Odor and taste not distinctive.

Basidiospores 10–13× 10–14.5 µm (11.8 ± 0.9 × 12.4 ± 1.0; range, mean ± SD), Q_m_ = 1.05, 6–8 angled with irregular pronounced angles, thick-walled, heterodiametrical. Basidia clavate, four-spored, 12–17× 37–52 µm. Cheilocystidia abundant, 10–16× 28–62 µm, cylindrical to narrowly clavate, thin-walled, contents granular. Pleurocystidia absent. Pileipellis a cutis composed of cylindrical to subcylindrical hyphae, almost hyaline, thin-walled, 5–14 µm wide, with scattered bundles of ascending hyphae. Pileitrama regular, made up of cylindrical hyphae, intracellular brown to reddish pigment, thin-walled, 5–12 µm wide. Clamp connections present in all hyphal tissues.

##### Additional specimens examined

J Sasari-Toge Pass, Kyoto, Kyoto Pref., Japan, on soil in a mixed forest of *Fagus crenata* and *Q. crispula* var. *crispula*, 16 Sep. 2020, voucher **(TNS-F-82985**), GenBank accession: LC786653 (ITS1); Tanakami, Otsu, Shiga Pref., Japan, on soil in a mixed forest of *Q. serrata* and *P. densiflora*, 28 Aug. 2020, voucher (**TNS-F-82986**), GenBank accession: LC786654 (ITS1), LC786705 (ITS2); 12 Sep. 2019, voucher (**TNS-F-82999**), GenBank accession: LC786667 (ITS1), LC786718 (ITS2); voucher (**TNS-F-83000**), GenBank accession: LC786668 (ITS1), LC786719 (ITS2); voucher (**TNS-F-83001**), GenBank accession: LC786669 (ITS1), LC786720 (ITS2); voucher (**TNS-F-83002**), GenBank accession: LC786670 (ITS1), LC786721 (ITS2); voucher (**TNS-F-83003**), GenBank accession: LC786671 (ITS1), LC786722 (ITS2); voucher **(TNS-F-83005**), GenBank accession: LC786673 (ITS1), LC786724 (ITS2); voucher (**TNS-F-83006**), GenBank accession: LC786674 (ITS1), LC786725 (ITS2); voucher (**TNS-F-83011**); 30 Sep. 2019, voucher (**TNS-F-83013**), GenBank accession: LC786676 (ITS1), LC786728 (ITS2); voucher (**TNS-F-83015**), GenBank accession: LC786678 (ITS1), LC786730 (ITS2); 11 Oct, 2019, voucher (**TNS-F-83016**), GenBank accession: LC786679 (ITS1), LC786731 (ITS2); voucher (**TNS-F-83022**), GenBank accession: LC786683 (ITS1), LC786737 (ITS2); Yakushima, Kagoshima Pref., Japan, on soil in a mixed forest of *Abies firma* and *Tsuga sieboldii*, 12 Sep. 2008, voucher (**KYO-YK125**), GenBank accession: LC786690 (ITS1), LC786743 (ITS2); Kutashimonocho, Sakyo-ku, Kyoto, Kyoto Pref., Japan, on soil in a mixed forest of *Q. crispula* var. *crispula* and *Castanea crenata*, 17 Sep. 2018, voucher (**KYO-HC59**), GenBank accession: LC786696 (ITS1), LC786749 (ITS2).

##### Distribution

Honshu and Kyushu, Japan.

##### Habitat

Solitary or gregarious on the ground in a mixed broadleaf-conifer forest from late August to mid-October. Especially abundant in forests where soil parent rocks are granite. Putatively soil decomposing fungi.

##### Remarks

This species has been confused with *E. quadratum* for a long time, but *E. quadratum* is distinguished by an orange-yellow to orange salmon pileus and smaller basidiospores (7–10 μm) [11]. This species is also somewhat confused with *E. flavescens*, which sometimes has tan to orange-tan basidiocarps. However, *E. flavescens* is characterized by paler basidiocarps, a shin-to-silky pileus and a pileipellis without or almost without bundles of ascending hyphae. The basidiospores of *E. flavescens* are slightly smaller than those of this species (Figs. 4–6), but the difference is not large enough to use as a diagnostic feature. Moreover, this species is somewhat morphological similar to *E. kovalenkoi*, species reported from Viet Nam, but *E. kovalenkoi* is characterized by bright orange basidiocarps and pileus without distinct papilla [43]. This species seems to prefer nutrient poor soils, such as granite-based soil, and has never been reported outside Japan. Thus, this species may have a restricted geographical distribution.

#### *Entoloma flavescens* Hirot. Sato, sp. nov. (Figs. 3D–F, 5)

##### MycoBank no

MB 10015984.

##### Holotype

Yoshidayama, Sakyo-ku, Kyoto, Kyoto Pref., Japan, on soil in a mixed forest of *Q. serrata*, *Q. glauca* and *Castanopsis cuspidata*, 29 Jul. 2019, voucher (**TNS-F-82995**), GenBank accessions: LC786663 (ITS1), LC786714 (ITS2)

##### Etymology

*flavescens*, Latin, yellow color (typical color of fresh basidiocarps)

##### Diagnosis

*Entoloma flavescens* sp. nov. is characterized by the following characteristics, a lemon-yellow to tan and shiny to silky pileus, basidiospores 8–13× 10–13 µm, pileipellis a cutis composed of cylindrical to subcylindrical hyphae without scattered bundles of ascending hyphae, and clamp connections in all hyphal tissues.

##### Description

Pileus 1–6 cm diam, thin, conical to campanulate, with an acute cusp at the apex; surface lemon-yellow (C 10, M 0, Y 60, K 0) to mustard yellow (C 10, M 35, Y 90, K 0), fading to tan (C 20, M 50, Y 80, K 0) or orange-tan (C 25, M 60, Y 80, K 0), dry, shiny when fresh, longitudinally striate, covered partially with whitish silky scales specifically near the apex, margin slightly lobed. Lamellae adnate to adnexed, subdistant, subventricose, margin serrate-fimbriate, concolorous with pileus, turning pinkish when mature. Flesh thin, fragile, concolorous with pileus. Stipe 4–13 cm long, 2–4 mm thick, slender, equal or subbulbous, striate fibrillose, often twisted, hollow, concolorous with pileus or paler, base white-mycelioid. Odor and taste not distinctive.

Basidiospores 9–13× 10–13 µm (11.0 ± 0.9 × 11.4 ± 0.8; range, mean ± SD), Q_m_ = 1.04, 6–8 angled with irregular pronounced angles, thick-walled, heterodiametrical. Basidia clavate, four-spored, 11–16× 31–52 µm. Cheilocystidia abundant, 10–16× 28–50 µm, cylindrical to narrowly clavate, thin-walled, contents granular. Pleurocystidia absent. Pileipellis a cutis, composing of cylindrical to subcylindrical hyphae, almost hyaline, thin-walled, 5–13 µm wide. Pileitrama regular, made up of cylindrical hyphae, intracellular brownish pigment, thin-walled, 5–13 µm wide. Clamp connections present in all hyphal tissues.

##### Additional specimens examined

Gero, Gif Pref., Japan, on soil in a mixed forest of *A. firma* and *T. sieboldii*, 09 Aug. 2019, voucher (**TNS-F-82996**), GenBank accessions: LC786664 (ITS1), LC786715 (ITS2); 22 Sep. 2018, voucher (**KYO-KS97**), GenBank accessions: LC786693 (ITS1); Mt. Maya, Hyogo Pref., Japan, 13 Jul. 2021, on soil in a mixed broadleaf forest, voucher (**TNS-F-82990**), GenBank accessions: LC786658 (ITS1), LC786709 (ITS2); Yakushima, Kagoshima Pref., Japan, on soil in a mixed forest of *A. firma* and *T. sieboldii*, 10 Sep. 2008, voucher (**KYO-YK71**), GenBank accessions: LC786688 (ITS1); 10 Sep. 2008, voucher (**KYO-YK77**), GenBank accessions: LC786689 (ITS1), LC786742 (ITS2); on soil in a mixed forest of *A. firma* and *Castanopsis sieboldii*, 23 Sep. 2008, voucher (**KYO-YK278**), GenBank accessions: LC786691 (ITS1), LC786744 (ITS2); 5 Oct. 2004, voucher (**KYO-YK489**), GenBank accessions: LC786692 (ITS1), LC786745 (ITS2); Kutashimonocho, Sakyo-ku, Kyoto, Kyoto Pref., Japan, on soil in a mixed forest of *Q. crispula* var. *crispula* and *C. crenata*, 17 Sep. 2018, voucher (**KYO-HC43**), GenBank accessions: LC786694 (ITS1), LC786747 (ITS2); Mt. Daimonji, Sakyo-ku, Kyoto, Kyoto Pref., Japan, on soil in a mixed forest of *Q. serrata* and *Q. glauca*, 31 Jul. 2020, voucher (**TNS-F-82980**), GenBank accessions: LC786648 (ITS1), LC786700 (ITS2); 31 Jul. 2020, voucher (**TNS-F-82983**), GenBank accessions: LC786651 (ITS1), LC786703 (ITS2); 31 Jul. 2020, voucher (**TNS-F-82984**), GenBank accessions: LC786652 (ITS1), LC786704 (ITS2); 06 Sep. 2019, voucher (**TNS-F-82997**), GenBank accessions: LC786665 (ITS1), LC786716 (ITS2); Yoshidayama, Sakyo-ku, Kyoto, Kyoto Pref., Japan, 15 Jul. 2019, voucher (**TNS-F-82977**), GenBank accessions: LC786645 (ITS1), LC786697 (ITS2); Tanakami, Otsu, Shiga Pref., Japan, on soil in a mixed forest of *Q. serrata* and *P. densiflora*, 28 Aug. 2020, voucher (**TNS-F-82987**), GenBank accessions: LC786655 (ITS1), LC786706 (ITS2); 28 Aug. 2020, voucher (**TNS-F-82988**), GenBank accessions: LC786656 (ITS1), LC786707 (ITS2); 12 Sep. 2019, voucher (**TNS-F-82998**), GenBank accessions: LC786666 (ITS1), LC786717 (ITS2); 11 Oct. 2019, voucher (**TNS-F-83017**), GenBank accessions: LC786680 (ITS1), LC786732 (ITS2); 02 Nov. 2019, voucher (**TNS-F-83025**), GenBank accessions: LC786686 (ITS1), LC786740 (ITS2); 02 Nov. 2019, voucher (**TNS-F-83026**), GenBank accessions: LC786687 (ITS1), LC786741 (ITS2).

##### Distribution

Japan (Honshu and Kyushu), China and Far Eastern Russia.

##### Habitat

Solitary or gregarious on the ground in a mixed broadleaf-conifer forest, from mid-July to early November. Putatively soil decomposing fungi.

##### Remarks

This species has been long confused with *E. murrayi*; however, *E. murrayi* is distinguished by smaller basidiospores (7–9.5 μm) and rare clamp connections [11]. This species sometimes has tan to orange-tan basidiocarps, somewhat similar to those of *E. quadratum*. However, the pileus of *E. quadratum* is not shiny or silky, and basidiospores of *E. quadratum* are smaller than those of this species (7–10 μm) [11]. In Japan, *E. album* was treated as the forma of this species due to the paucity of diagnostic characters [44]. However, the basidiocarps of *E. album* never become yellowish except for papilla. Moreover, *E. album* is characterized by thinner cheilocystidia (Figs. 5, 6) and thinner pileipellis hyphae. This species can be also distinguished from *E. luteum* [45] and *E.* aff. *luteum* in Japan by having an acute cusp at the apex, more vivid pileus color, and much shorter cheilocystidia.

#### Entoloma album Hiroë, [46] Fig. 3G–I

##### MycoBank no

MB 250576.

##### Holotype

Inaba, Tottori Pref., Japan (the type material cannot be traced in the mycological herbaria in Japan, including TNS and the Tottori University mycological herbarium (TUMH), and is probably lost).

##### Description

Pileus 1–4 cm diam, thin, conical to campanulate, with an acute cusp at the apex; surface white (C 0, M 0, Y 0, K 0), pale white (C 10, M 2, Y 10, K 0) to cream white (C 10, M 10, Y 20, K 0), dry, shiny when fresh, longitudinally striate, covered partially with whitish silky scales specifically near the apex, margin slightly lobed. Lamellae adnate to adnexed, subdistant, subventricose, margin serrate-fimbriate, concolorous with pileus, turning pinkish when mature. Flesh thin, fragile, concolorous with pileus. Stipe 3–10 cm long, 2–4 mm thick, slender, equal or subbulbous, striate fibrillose, often twisted, hollow, concolorous with pileus or paler, base white-mycelioid. Odor and taste not distinctive.

Basidiospores 9–13× 10–13 µm (10.7 ± 0.9 × 11.3 ± 0.8; range, mean ± SD), Q_m_ = 1.06, 6–8 angled with irregular pronounced angles, thick-walled, heterodiametrical. Basidia clavate, four-spored, 10–15× 38–59 µm. Cheilocystidia abundant, 8–12× 32–52 µm, cylindrical to narrowly clavate, thin-walled, contents granular. Pleurocystidia absent. Pileipellis a cutis, composing of cylindrical to subcylindrical hyphae, almost hyaline, thin-walled, 3–6 µm wide. Pileitrama regular, made up of cylindrical hyphae, almost hyaline, thin-walled, 3–6 µm wide. Clamp connections present in all hyphal tissues.

##### Specimens examined

Yoshidayama, Sakyo-ku, Kyoto, Kyoto Pref., Japan, on soil in a mixed forest of *Q. serrata*, *Q. glauca* and *C. cuspidata*, 18 Jul. 2019, voucher (**TNS-F-82991**), GenBank accessions: LC786659 (ITS1), LC786710 (ITS2); Mt. Maya, Hyogo Pref., Japan, on soil in a mixed broadleaf forest, 20 Jul. 2019, voucher **(TNS-F-82992**), GenBank accessions: LC786660 (ITS1), LC786711 (ITS2); 20 Jul. 2019, voucher (**TNS-F-82994**), GenBank accessions: LC786662 (ITS1), LC786713 (ITS2); Tanakami, Otsu, Shiga Pref., Japan, on soil in mixed forest of *Q. serrata* and *P. densiflora*, 12 Sep. 2019, voucher (**TNS-F-83004**), GenBank accessions: LC786672 (ITS1), LC786723 (ITS2); 11 Oct. 2019, voucher (**TNS-F-83018**), GenBank accessions: LC786681 (ITS1), LC786733 (ITS2); 02 Nov. 2019, voucher (**TNS-F-83023**), GenBank accessions: LC786684 (ITS1), LC786738 (ITS2); 02 Nov. 2019, voucher (**TNS-F-83024**), GenBank accessions: LC786685 (ITS1), LC786739 (ITS2); Kutashimonocho, Sakyo-ku, Kyoto, Kyoto Pref., Japan, on soil in mixed forest of *Q. crispula* var. *crispula* and *C. crenata*, 17 Sep. 2018, voucher (**KYO-HC41**), GenBank accessions: LC786695 (ITS1), LC786748 (ITS2).

##### Distribution

Japan (Honshu) and China.

##### Habitat

Solitary or gregarious on the ground in mixed forests of broadleaf and coniferous trees, from mid-July to early November. Putatively soil decomposing fungi.

##### Remarks

This species was previously treated as the forma of *E. murrayi* in Japan [44], but these two species should be treated as independent species. This species can be distinguished from both of *E. murrayi* and *E. flavescens* in having white basidiomes and thinner cheilocystidia. Whitish basidiocarps of this species are somewhat similar with those of *E. cycneum* and *E. peristerinum*, both of which were reported from Viet Nam [47]. However, these two species are characterized by clearly hygrophanous pileus, smaller basidiospores (*E. cycneum*: 8.0–8.5 × 8.5–9.0 μm, *E. peristerinum*: 7–8 × 8–9.5 μm) and longer cheilocystidia (*E. cycneum*: 95–160 × 7.5–9 μm, *E. peristerinum*: 75– 215 × 12–15 μm) [47].

## Discussion

Results of molecular phylogeny and population genetics using the nuc_concat dataset indicate that *E. album*, *E. flavescens* and *E. kermesinum* represent distinct species. The molecular phylogeny of the nuc_concat dataset showed that genetic variations among *E. album*, *E. flavescens* and *E. kermesinum* were much larger than those within respective species (Fig. 2), suggesting restricted gene flow among the three species. Nevertheless, molecular phylogenies indicated that *E. album* and *E. flavescens* were closely related with small genetic differentiation. Indeed, cytonuclear disequilibrium —a strong evidence for restricted gene flow [48, 49] —was not observed between *E. album* and *E. flavescens*, probably due to the incomplete lineage sorting of mtLSU. However, results of AMOVA using the nuc_concat dataset showed that gene flow was strongly restricted between *E. album* and *E. flavescens* even in areas where two species coexisted (Table 1), suggesting that these species are reproductively isolated from each other. Consequently, our results support that *E. album*, *E. flavescens* and *E. kermesinum* should be treated as independent species.

The molecular phylogeny of the ITS dataset also has important implications for the taxonomy of the *E. quadratum*–*murrayi* complex. The clades of *E. quadratum* and *E. murrayi* were phylogenetically distinct from the clades of *E. kermesinum* and *E. flavescens*, respectively (Fig. 1B), supporting the hypothesis that these species are not conspecific. Notably, the molecular phylogenetic tree of the ITS dataset showed that the *E. quadratum* and *E. murrayi* clades comprised specimens collected near type localities (i.e., Eastern North America), suggesting that they represent the focal species. Moreover, *E. kermesinum* was phylogenetically distinct from *E. kovalenkoi*, which is somewhat morphologically similar to *E. kermesinum* [43] (Fig. 1B). Similarly, *E. album* was phylogenetically distant from both of *E. cycneum* and *E. peristerinum*, both of which are characterized by whitish basidiocarps [47] (Fig. 1B). These findings indicate that *E. album*, *E. flavescens* and *E. kermesinum* represent independent species.

This study demonstrates the diagnostic morphological features of *E. kermesinum, E. flavescens* and *E. album*. For instance, *E. kermesinum* can be distinguished from *E. flavescens* and *E. album* by crimson to deep salmon red basidiocarps and a fibrillose pileus with whitish fibrous scales (Fig. 3). Notably, *E. flavescens* sometimes has a tan to orange-tan basidiocarp, and thus is somewhat confused with *E. kermesinum* (Fig. 3F). Specimens with tan to orange-tan basidiocarps were not closely related (Figs. 1B, 2), suggesting that this characteristic is not heritable (e.g., commonly observed in old basidiocarps). Considering such color variations of basidiocarps, *E. flavescens* is easily distinguished from *E. kermesinum* because *E. flavescens* has a shiny-to-silky pileus and a pileipellis without or almost without bundles of ascending hyphae. In addition, *E. album* is somewhat confused with *E. flavescens* [44], but *E. album* can be distinguished by whitish basidiocarps (Fig. 3). The basidiocarps of *E. album* are never yellowish, except for the papilla. Moreover, *E. album* is distinguished from *E. flavescens* by thinner cheilocystidia (Figs. 5, 6). Overall, macroscopic and microscopic morphological features of the pileus surface are especially useful for distinguishing *E. album*, *E. flavescens*, and *E. kermesinum*.

Morphological characteristics of *E. kermesinum*, *E. flavescens,* and *E. album* are distinguishable from those of related species whose type locality is outside Japan. Basidiocarp color of *E. kermesinum* (crimson to deep red) is much deeper than those of *E. quadratum* (orange-yellow to orange salmon) [11] and *E. kovalenkoi* (bright orange) [43]. In addition, relatively large basidiospores and pileus with distinct papilla are diagnostic features distinguishing *E. kermesinum* from *E. quadratum* and *E. kovalenkoi*, respectively [11, 43]. Similarly, basidiospores of *E. flavescens* are larger than those of *E. murrayi* [11]. *Entoloma flavescens* is also characterized by clamp connections, which are absent or rare in *E. murrayi* [11]. Moreover, *E. album* is distinguished from both of *E. cycneum* and *E. peristerinum* by having larger basidiospores and shorter cheilocystidia [47]. These morphological characteristics are useful for distinguishing *E. kermesinum*, *E. flavescens,* and *E. album* from other species.

Ecological and biogeographic features of *E. kermesinum* appear to be distinct from those of other species. Most *E. kermesinum* basidiocarp were found in Tanakami (Otsu, Shiga Pref., Japan), where soil parent rocks are granite, and none were found outside Japan. By contrast, *E. flavescens* and *E. album* were found in more diverse areas, including areas outside Japan (Figs. 1, 2). Thus, *E. kermesinum* may prefer nutrient poor soils and have restricted geographic distribution. Although the basidiocarps of *E. kermesinum* and *E. flavescens* are believed to grow together frequently [28], this is presumably not true. It seems more likely that *E. flavescens* with tan to orange-tan basidiocarps was misidentified as *E. kermesinum*. Notably, intraspecific variations in *E. kermesinum* were much smaller than those in *E. album* and *E. flavescens*, even when restricted to specimens collected in Tanakami (Figs. 1, 2), suggesting presence of a bottleneck effect or founder effect in *E. kermesinum* populations. Thus, *E. kermesinum* has a distinctive ecological habitat and biogeography.

This study provides useful information about the taxonomy of species related to the *E. quadratum*–*murrayi* complex. Based on molecular phylogenetic trees, one specimen of *E.* sp. (EN08) was found to be closely related to, but distinguished from, *E. album* and *E. flavescens* (Figs. 1, 2). Nevertheless, only one specimen is available for this hypothetical species, and more samples are required to clarify intraspecific variations and quantify gene flow. Moreover, *E.* aff. *luteum* in East Asian countries (i.e., Japan and China) were phylogenetically distinct from *E. luteum* collected in Canada, near the type locality, i.e., New York [45], suggesting that East Asian materials that had been identified as *E. luteum* may be distinct species. Further sampling and examination are needed to support this hypothesis.

## Acknowledgements

We thank Dr. Hiroki Yamanaka (Center for Biodiversity Science, Ryukoku University) for supporting the laboratory work, especially the high-throughput sequencing using the Illumina MiSeq. We also thank Dr. Toshimitsu Fukiharu (Natural History Museum and Institute, Chiba), who helped the field survey of this study. We are grateful to Dr. Kasuya Taiga (Keio University) for useful comments for this study.

## Author contributions

H.S. planned and designed the research, performed the field surveys, the experiments, analyzed the data, interpreted the results, and wrote the manuscript. O.S. performed the field surveys, the experiments, analyzed the data, interpreted the results, and wrote the manuscript. Y.S. performed the field surveys, the experiments, and interpreted the results. H.S. and O.S. contributed equally and should be considered as co-first authors.

## Legends of supplementary materials

**S1 Table. Sample lists used in this study.** GenBank accession number are provided for the consensus sequences of the nuclear ribosomal internal transcribed spacer 1 (ITS1) and spacer 2 (ITS2) regions (NA: not available).

**S2 Table. List of PCR primers used in this study.** Forward and reverse primers are fused with Illumina sequencing primer regions and 6-mer Ns (5’-TCGTCGGCAGCGTCAGATGTGTATAAGAGACAGNNNNNN [forward primer]-3’ and 5’-GTCTCGTGGGCTCGGAGATGTGTATAAGAGACAGNNNNNN [reverse primer]-3’).

## References

1. Cao B, Haelewaters D, Schoutteten N, Begerow D, Boekhout T, Giachini AJ, et al. Delimiting species in Basidiomycota: a review. Fungal Diversity. 2021:1–57. doi: 10.1007/s13225-021-00479-5.

2. Sato H, Ohta R, Murakami N. Molecular prospecting for cryptic species of the *Hypholoma fasciculare* complex: toward the effective and practical delimitation of cryptic macrofungal species. Scientific Reports. 2020;10(1):13224. doi: 10.1038/s41598-020-70166-z.

3. Sato H. The evolution of ectomycorrhizal symbiosis in the Late Cretaceous is a key driver of explosive diversification in Agaricomycetes. New Phytologist. 2023. doi: 10.1111/nph.19055.

4. Nilsson RH, Kristiansson E, Ryberg M, Hallenberg N, Larsson KH. Intraspecific ITS variability in the kingdom Fungi as expressed in the international sequence databases and its implications for molecular species identification. Evolutionary bioinformatics online. 2008;4:193. doi: 10.4137/EBO.S653.

5. Schoch CL, Seifert KA, Huhndorf S, Robert V, Spouge JL, Levesque CA, et al. Nuclear ribosomal internal transcribed spacer (ITS) region as a universal DNA barcode marker for Fungi. Proceedings of the National Academy of Sciences. 2012;109(16):6241–6. doi: 10.1073/pnas.1117018109.

6. Koufopanou V, Burt A, Taylor JW. Concordance of gene genealogies reveals reproductive isolation in the pathogenic fungus *Coccidioides immitis*. Proceedings of the National Academy of Sciences of the United States of America. 1997;94(10):5478. doi: 10.1073/pnas.94.10.5478.

7. Sato H, Yumoto T, Murakami N. Cryptic species and host specificity in the ectomycorrhizal genus *Strobilomyces* (Strobilomycetaceae). American Journal of Botany. 2007;94(10):1630–41. doi: 10.3732/ajb.94.10.1630.

8. Taylor JW, Jacobson DJ, Kroken S, Kasuga T, Geiser DM, Hibbett DS, Fisher MC. Phylogenetic species recognition and species concepts in fungi. Fungal Genetics and Biology. 2000;31(1):21–32. doi: 10.1006/fgbi.2000.1228. PubMed PMID: ISI:000166290700003.

9. Kobmoo N, Mongkolsamrit S, Arnamnart N, Luangsa-ard JJ, Giraud T. Population genomics revealed cryptic species within host-specific zombie-ant fungi (*Ophiocordyceps unilateralis*). Molecular phylogenetics and evolution. 2019;140:106580. doi: 10.1016/j.ympev.2019.106580.

10. Sato H, Murakami N. Reproductive isolation among cryptic species in the ectomycorrhizal genus *Strobilomyces*: Population-level CAPS marker-based genetic analysis. Molecular Phylogenetics and Evolution. 2008;48(1):326–34. doi: 10.1016/j.ympev.2008.01.033.

11. Horak E. On cuboid-spored species of *Entoloma* (Agaricales). Sydowia. 1976;28:171–236.

12. Horak E. Fungi of New Zealand: Pluteaceae (Pluteus, Volvariella), Entolomataceae (Claudopus, Clitopilus, Entoloma, Ponzarella, Rhodocybe, Richoniella): Fungal Diversity Press; 2008.

13. Morozova O, Noordeloos ME, Vila J. *Entoloma* subgenus *Leptonia* in boreal-temperate Eurasia: towards a phylogenetic species concept. Persoonia-Molecular Phylogeny and Evolution of Fungi. 2014;32(1):141–69. doi: 10.3767/003158514X681774.

14. Morozova O, Popov E, Alexandrova A, Pham THG, Noordeloos ME. Four new species of *Entoloma* (Entolomataceae, Agaricomycetes) subgenera *Cyanula* and *Claudopus* from Vietnam and their phylogenetic position. Phytotaxa. 2022;549(1):1–21. doi: 10.11646/phytotaxa.549.1.1.

15. Morozova O, Popov E, Kovalenko A. Studies on mycobiota of Vietnam. I. Genus *Entoloma*: New records and new species. Mikologiya i fitopatologiya. 2012;46(3):182–200.

16. Noordeloos ME, Gates GM, Sachs AW. The Entolomataceae of Tasmania. Dordrecht, Netherlands: Springer; 2012.

17. Noordeloos ME, Hausknecht A. The genus *Entoloma* (Basidiomycetes, Agaricales) of the Mascarenes and Seychelles. Fungal Diversity. 2007;27(1):111–44.

18. Noordeloos ME, Morozova OV. New and noteworthy *Entoloma* species from the Primorsky Territory, Russian Far East. Mycotaxon. 2010;112:231–55. doi: 10.5248/112.231.

19. Reschke K, Noordeloos ME, Manz C, Hofmann TA, Rodríguez-Cedeño J, Dima B, Piepenbring M. Fungal diversity in the tropics: *Entoloma* spp. in Panama. Mycological Progress. 2022;21(1):93–145. doi: 10.1007/s11557-021-01752-2.

20. Noordeloos ME. Introduction to the taxonomy of the genus *Entoloma* sensu lato (Agaricales). Persoonia-Molecular Phylogeny and Evolution of Fungi. 1981;11(2):121–51.

21. Singer R. The Agaricales in modern taxonomy. 4th edn ed. Koenigstein: Koeltz Scientific Books; 1986.

22. Karstedt F, Capelari M, Baroni TJ, Largent DL, Bergemann SE. Phylogenetic and morphological analyses of species of the Entolomataceae (Agaricales, Basidiomycota) with cuboid basidiospores. Phytotaxa. 2019;391(1):1–27-1–. doi: 10.11646/phytotaxa.391.1.1.

23. Morgado L, Noordeloos ME, Lamoureux Y, Geml J. Multi-gene phylogenetic analyses reveal species limits, phylogeographic patterns, and evolutionary histories of key morphological traits in *Entoloma* (Agaricales, Basidiomycota). Persoonia-Molecular Phylogeny and Evolution of Fungi. 2013;31(1):159–78. doi: 10.3767/003158513X673521.

24. Põlme S, Abarenkov K, Nilsson RH, Lindahl BD, Clemmensen KE, Kauserud H, et al. FungalTraits: a user-friendly traits database of fungi and fungus-like stramenopiles. Fungal diversity. 2020;105(1):1–16. doi: 10.1007/s13225-020-00466-2.

25. Tedersoo L, May TW, Smith ME. Ectomycorrhizal lifestyle in fungi: global diversity, distribution, and evolution of phylogenetic lineages. Mycorrhiza. 2010;20(4):217–63. doi: 10.1007/s00572-009-0274-x.

26. Hobbie EA, Weber NS, Trappe JM. Mycorrhizal vs saprotrophic status of fungi: the isotopic evidence. New Phytologist. 2001;150(3):601–10. doi: 10.1046/j.1469-8137.2001.00134.x.

27. Kobayashi H, Hatano K. A morphological study of the mycorrhiza of *Entoloma clypeatum* f. *hybridum* on *Rosa multiflora*. Mycoscience. 2001;42:83–90. doi: 10.1007/BF02463979.

28. Imazeki R, Hongo T. Colored illustrations of mushrooms of Japan. Osaka: Hoikusha; 1987.

29. Sato H, Tsujino R, Kurita K, Yokoyama K, Agata K. Modelling the global distribution of fungal species: new insights into microbial cosmopolitanism. Molecular Ecology. 2012;21:5599–612. doi: 10.1111/mec.12053.

30. Sato H, Tanabe AS, Toju H. Host shifts enhance diversification of ectomycorrhizal fungi: diversification rate analysis of the ectomycorrhizal fungal genera *Strobilomyces* and *Afroboletus* with an 80-gene phylogeny. New Phytologist. 2017;214(1):443–54. doi: 10.1111/nph.14368.

31. Toju H, Tanabe AS, Yamamoto S, Sato H. High-Coverage ITS Primers for the DNA-Based Identification of Ascomycetes and Basidiomycetes in Environmental Samples. PloS one. 2012;7(7):e40863. doi: 10.1371/journal.pone.0040863.

32. Hamady M, Walker JJ, Harris JK, Gold NJ, Knight R. Error-correcting barcoded primers for pyrosequencing hundreds of samples in multiplex. Nature methods. 2008;5(3):235–7. doi: 10.1038/nmeth.1184.

33. Tanabe AS, Toju H. Two new computational methods for universal DNA barcoding: A benchmark using barcode sequences of bacteria, archaea, animals, fungi, and land plants. PloS one. 2013;8(10):e76910. doi: 10.1371/journal.pone.0076910.

34. Rognes T, Flouri T, Nichols B, Quince C, Mahé F. VSEARCH: a versatile open source tool for metagenomics. PeerJ. 2016;4:e2584. doi: 10.7717/peerj.2584.

35. Edgar RC, Haas BJ, Clemente JC, Quince C, Knight R. UCHIME improves sensitivity and speed of chimera detection. Bioinformatics. 2011;27(16):2194–200. doi: 10.1093/bioinformatics/btr381.

36. R Core Team. R: A Language and Environment for Statistical Computing. R Foundation for Statistical Computing, Vienna, Austria. <https://www.R-project.org/>. 2023.

37. Katoh K, Misawa K, Kuma K, Miyata T. MAFFT: a novel method for rapid multiple sequence alignment based on fast Fourier transform. Nucleic acids research. 2002;30(14):3059–66. doi: 10.1093/nar/gkf436.

38. Charif D, Lobry JR. SeqinR 1.0-2: a contributed package to the R project for statistical computing devoted to biological sequences retrieval and analysis. In: Bastolla U, Porto M, Roman HE, Vendruscolo M, editors. Structural approaches to sequence evolution. Berlin: Springer; 2007. p. 207–32.

39. Stamatakis A. RAxML-VI-HPC: maximum likelihood-based phylogenetic analyses with thousands of taxa and mixed models. Bioinformatics. 2006;22(21):2688–90. doi: 10.1093/bioinformatics/btl446.

40. Tanabe AS. Kakusan4 and Aminosan: two programs for comparing nonpartitioned, proportional and separate models for combined molecular phylogenetic analyses of multilocus sequence data. Molecular Ecology Resources. 2011;11(5):914–21. doi: 10.1111/j.1755-0998.2011.03021.x.

41. Kamvar ZN, Tabima JF, Grünwald NJ. Poppr: an R package for genetic analysis of populations with clonal, partially clonal, and/or sexual reproduction. PeerJ. 2014;2:e281. doi: 10.7717/peerj.281.

42. Paradis E. pegas: an R package for population genetics with an integrated–modular approach. Bioinformatics. 2010;26(3):419–20. doi: 10.1093/bioinformatics/btp696.

43. Crous PW, Osieck ER, Jurjević Ž, Boers J, Van Iperen A, Starink-Willemse M, et al. Fungal Planet description sheets: 1284–1382. Persoonia: Molecular Phylogeny and Evolution of Fungi. 2021;47:178. doi: 10.3767/persoonia.2022.48.08.

44. Hongo T. Notes on Japanese larger fungi (1). The Journal of Japanese Botany. 1951;26(1):23–6.

45. Peck CH. Report of the State Botanist (1900). Annual report on the New York State Museum of Natural History. 1902;54:131–99.

46. Hiroe I. *Entoloma* of Japan (2). Notes on larger fungi of the San-in district. (VII.). Applied Mushroom Science. 1939;4(1):1–3.

47. Morozova O, Pham THG. New species of *Entoloma* subgenera *Cubospora* and *Leptonia* (Agaricales, Basidiomycota) from central Vietnam. Journal of Fungi. 2023;9(6):621. doi: 10.3390/jof9060621.

48. Arnold J. Cytonuclear Disequilibria in Hybrid Zones. Annu Rev Ecol Syst. 1993;24:521–54. doi: 10.1146/annurev.es.24.110193.002513. PubMed PMID: ISI:A1993MJ37100019.

49. Avise JC. Phylogeography: the history and formation of species: Harvard university press; 2000.

